# Storage root yield of sweetpotato as influenced by sweetpotato leaf curl virus and its interaction with sweetpotato feathery mottle virus, and sweetpotato chlorotic stunt virus in Kenya

**DOI:** 10.1101/795518

**Authors:** Bramwel W. Wanjala, Elijah M. Ateka, Douglas W. Miano, Jan W. Low, Jan F. Kreuze

**Affiliations:** International Potato Center, SSA Regional Office, PO Box 25171 - 00603, Nairobi, Kenya.; Jomo Kenyatta University of Agriculture and Technology, P.O. Box 62000 – 00200, Nairobi, Kenya.; International Potato Center, Avenida La Molina 1895, La Molina. Apartado Postal 1558, Lima, Peru.; The University of Nairobi, P.O. Box: 30197, 00100 Nairobi, Kenya.

**Keywords:** SPLCV, sweepovirus, SPFMV, SPCSV, treatment, yield

## Abstract

The effect of a Kenyan strain of sweetpotato leaf curl virus (SPLCV) and its interactions with sweetpotato feathery mottle virus (SPFMV), and sweetpotato chlorotic stunt virus (SPCSV) on root yield was determined. Trials were performed during two seasons using varieties contrasting in their resistance to sweetpotato virus disease, ‘Kakamega’ and ‘Ejumula’, in a randomized complete block design with sixteen treatments replicated three times. The treatments included plants graft inoculated with SPLCV, SPFMV and SPCSV alone and in possible dual or triple combinations. Yield and yield related parameters were evaluated at harvest. Results showed marked differences in the effect of SPLCV infection on the two varieties: ‘Ejumula’, which is susceptible to SPFMV and SPCSV, suffered no significant yield loss from SPLCV infection, whereas ‘Kakamega’, which is more resistant to SPFMV and SPCSV, suffered an average of 47% yield loss, despite only mild symptoms occurring in both varieties. These results highlight the variability in sensitivity to SPLCV between sweetpotato cultivars as well as a lack of correlation of SPLCV related symptoms with susceptibility to the virus. In addition, they underline the lack of correlation between resistance to the RNA viruses SPCSV and SPFMV and DNA virus SPLCV.

## Introduction

Ranked seventh in global food crop production, Sweetpotato (*Ipomoea batatas*) is the third most important root and tuber crop after potato and cassava. In the developing world, it ranks fourth in importance after rice, wheat, and corn (Kays, 2005). It is one of the traditional crops that play an important role in addressing food insecurity in most rural households in Africa (Gruneberg et al. 2015). Orange-fleshed sweetpotato varieties’ high β-carotene (source of pro-vitamin A) has seen an increased utilization in food and dietary programs aimed at addressing vitamin A deficiency; a global challenge in Sub-Saharan Africa (SSA) (Kurabachew, 2015). The crop is cultivated all-round the year, producing high yields under marginal conditions. Sweetpotato yields differ, from over 25 metric tons per hectare with high-input to below 3 metric tons per hectare when grown as a subsistence crop with minimal input (Ling et al. 2010). In Kenya, sweetpotato production is hindered by numerous biotic, abiotic and social factors (Kivuva et al. 2015). Pests and diseases are the greatest limitation that affect production and reduce yields (Motsa et al. 2015). Viral diseases are the greatest threat to sweetpotato production, causing yield losses of up to 80% (Gibson and Kreuze, 2015).

Recycling of vine cuttings leads to a significant decline in root yield and quality due to virus accumulation. Sweetpotato virus disease (SPVD), caused by dual infection of a Potyvirus, sweet potato feathery mottle virus (SPFMV), and a Crinivirus, sweetpotato chlorotic stunt virus (SPCSV), is most devastating in East Africa (Karyeija et al. 1998; Gibson et al. 1997). SPFMV is common in sweetpotato producing regions around the world (Ateka et al. 2004). SPCSV induces synergistic interactions with other sweetpotato viruses blonging to the species sweetpotato mild mottle virus (SPMMV) (Tairo et al. 2005), sweetpotato virus G (SPVG) (IsHak et al. 2003), and cucumber mosaic virus (CMV) (Cohen and Loebenstein, 1991). However, yield decline attributable to these viruses is cultivar dependent and previous studies have given contradictory findings. Milgram et al. (1996) and Clark and Hoy (2006), reported that single infection with SPFMV, SPVG, or isolates of the species sweetpotato virus 2 (syn. Ipomoea vein mosaic virus) did not considerably affect yield. In the contrary, Gutierrez et al. (2003) found that SPFMV-infected plants produced better yield than the healthy control. On the other hand, Gibson et al. (1997), Mukasa (2004), Njeru et al. (2004), Domola et al. (2008), reported yield reduction of up to 46%.

To date, over 30 viruses have been characterized as pathogens of sweetpotato, half of them belonging to the families *Geminiviridae* and *Caulimoviridae* (Clark et al. 2012). Members of the species sweet potato leaf curl virus (SPLCV) and related viruses infecting sweetpotato, belong to the genus *begomovirus* in the family *Geminiviridae*. They are highly variable making their taxonomy, which has been revised over recent years, problematic, but they can be distinguished from begomoviruses infecting other crops by their phylogenetically unique lineage, referred to as sweepoviruses (Albuquerque et al. 2012; Esterhuizen et al. 2012; Fauquet and Stanley, 2003; Wasswa et al. 2011, Cuellar et al. 2015). We will refer to them as such in this manuscript when discussing them in general, rather than individual isolates. Sweepoviruses are transmitted through vegetative propagation and semi-persistently by whiteflies (*Bemisia tabaci*). They have been isolated from sweetpotato fields in different parts of the world including the United States, South America, the Middle East, Southeast Asia, and East Africa (Briddon et al. 2006; Luan et al. 2006; Miano et al. 2006; Prasanth and Hegde 2008; Lozano et al. 2009; Paprotka et al. 2010; Albuquerque et al. 2012; Wasswa et al. 2011). Sweepovirus infected plants may exhibit upward curling and/or rolling of leaves, vein swelling, and vein mottle in young sweetpotato plants.

However, symptom remission is observed in mature plants, and most plants become symptomless (Miano et al. 2006). Sweepovirus single viral infections often lack obvious symptoms making it difficult to be recognized by growers. Miano et al.(2006) reported the occurrence of sweepoviruses in an agricultural field station in Kenya. Countrywide surveys conducted in 2011 (Maina, 2014; Maina et al. 2017) and (Wanjala, 2016/2017 - *unpublished data*) confirmed sweepoviruses to be present in the major sweetpotato growing regions of the country. The presence of sweepovirus inoculum in major sweetpotato producing areas in Kenya and the continuing expansion of the vector - *Bemisia tabaci* (Simmons et al. 2008) might have contributed to its broad geographic distribution in Kenya.

Despite the lack of characteristic foliar symptoms, sweepoviruses have been reported to cause between 10 and 80% yield loss for different sweet potato cultivars (Clark and Hoy 2006; Ling et al. 2010; Gibson and Kreuze, 2015). Studies have demonstrated that sweepoviruses when co-infected with SPCSV can lead to increased viral titres and symptoms in sweetpotatoes under controlled conditions (Cuellar et al. 2015). However, limited knowledge exists on the interaction of sweepoviruses, SPFMV and SPCSV in sweetpotato under field conditions or their effect on yield and quality of sweetpotato roots. Therefore, the study aimed at evaluating the effect of Kenyan isolates of a sweepovirus (SPLCV), SPFMV, and SPCSV alone, and co-infections on sweetpotato root yield of two cultivars contrasting in their resistance to SPVD.

## Methods and Materials

### Sources of healthy planting and detection of sweetpotato viruses

Clean virus tested (VT) *in vitro* planting materials were obtained from the International Potato Centre (CIP) germplasm collection at Kenya Plant Health Inspectorate Services - Plant Quarantine and Biosecurity Station (KEPHIS-PQBS) Muguga, Kenya. Plantlets of varieties ‘Kakamega’ and ‘Ejumula’ were hardened in insect proof greenhouses and away from plants that might be infected with viruses. Both cultivars are landraces, widely adaptable, have good storage root shapes if grown in light soils, high dry matter content, and excellent consumer acceptance, especially among children and women (Mwanga et al. 2007). Ejumula is susceptible to SPVD while Kakamega shows levels of field resistance to sweetpotato virus disease.

Biological indexing was carried out as described by Dennien et al. (2013) on *Ipomoea setosa* (indicator plant) that is highly sensitive to most sweetpotato infecting viruses. Vines singly infected with SPFMV, SPFMV and SPLCV were used as scions to an *Ipomoea setosa* stock seedling following the procedures in (Beetham and Mason 1992 and Dennien et al. 2013). Virus infection treatments (T1–T16) are described in Table 1. *I. setosa* seedling was grown out to 10 nodes (4-6 weeks after planting) and grafted with 2 two-node scions from the test plant, one from the basal portion of the vine and one from near the apex of the vine. A wedge graft was made at about 3 nodes above the cotyledonary node and a side veneer graft just below the cotyledonary node. Grafted plants in the pots were covered with plastic bags and placed into large, shallow trays lined with plastic sheeting. The *I. setosa* indicator plant was allowed to grow. To capture transient symptoms, indicator plants are observed twice weekly until 21 days post grafting (PG), then weekly until 42 days PG. The I. setosa was cut back above the graft site and allowed to regrow for an additional 3-4 weeks, continually observing for symptom development. Symptoms typical of different viruses as illustrated in Clark et al., 2012 and Dennien et al., 2013 were recorded.

**Table 1.** Description of treatments (viruses and their combinations) used to evaluate the effect of different viruses on sweetpotato varieties Ejumula and Kakamega.

| Treatment | Plot No | Cultivar | Treatment description |
| --- | --- | --- | --- |
| T1 | 15/30/43 | Ejumula | Non infected |
| T2 | 5/17<sup>1</sup>/40 |  | SPCSV |
| T3 | 16/31/47 |  | SPFMV |
| T4 | 9/27/41 |  | SPLCV +SPCSV |
| T5 | 3/22/33 |  | SPLCV +SPFMV |
| T6 | 12/23/46 |  | SPVD |
| T7 | 14/19/36 |  | SPLCV +SPVD |
| T8 | 10/25/44 |  | SPLCV |
| T9 | 4/24<sup>2</sup>/39 | Kakamega | Non infected |
| T10 | 11/29/34 |  | SPCSV |
| T11 | 1/20/48 |  | SPFMV |
| T12 | 7/21/35 |  | SPLCV +SPCSV |
| T13 | 6/32/42 |  | SPLCV +SPFMV |
| T14 | 8/26/38 |  | SPVD |
| T15 | 13/28/37 |  | SPLCV +SPVD |
| T16 | 2/18<sup>3</sup>/45 |  | SPLCV |

**Key:** SPCSV - *Sweet potato chlorotic stunt virus*, SPFMV - *Sweet potato feathery mottle virus* , SPLCV – *Sweet potato leaf curl virus* and SPVD - *Sweet potato virus disease*.
<sup>1</sup> Plot positive for sweepovirus in bulk PCR test at end of season I
<sup>2</sup> Plot positive for SPCSV in bulk PCR at end of season I
<sup>3</sup> Plot positive for SPFMV in bulk PCR test at end of season I, and SPCSV at end of season II

A standard Nitrocellulose membrane enzyme-linked immunosorbent assay (NCM-ELISA) was done using a test kit manufactured by the International Potato Center and as and described by Dennien et al (2013). It tests for 10 known sweetpotato infecting viruses: (C-6, CMV, SPCaLV, SPCV, SPCFV, SPCSV, SPFMV, SPLV, SPMMV, SPMSV and SPVG). It is a prerequisite for the test to use material that is first grafted onto I. setosa. This increases the virus concentration in the indicator and prevents inhibitors present in sweetpotato sap. There are no antisera available for SPLCV and sweepoviruses were tested by PCR as described by Li et al. (2004); using Sweepovirus-specific primers SPG1 (5’-CCC CKG TGC GWR AAT CCA T-3’) and SPG2 (5’-ATC CVA AYW TYC AGG GAG CTA A-3’), designed to amplify a 901-bp region encompassing partial AC1 and AC2 open reading frames (ORFs).

### Source of virus inoculum and virus inoculation

Plants singly infected with SPCSV (isolate KE_4) and SPFMV (isolate KE_42). used for graft infection were obtained from KEPHIS-PQBS. Sweepovirus (SPLCV) isolate KE_97 positive plants were collected in different parts of Kenya during surveillance surveys. Viruses were confirmed by grafting to *I. setosa* and use of NCM ELISA. In addition, the plants were subjected to screening by PCR for begomovirus as described above by Li et al.(2004). SPCSV and SPFMV were tested with Reverse Transcription PCR (RT-PCR) as described by Kwak et al. (2014). Furthermore, local strains of sweepovirus positive samples were confirmed by Sanger sequencing of the PCR product (GenBank id MN122257) and confirmed isolate KE_97 was a sweepovirus most closely related to SPLCV and we will refer to it as SPLCV from here onwards. Two-node cuttings were obtained from the VT hardened mother plants of ‘Kakamega’ and ‘Ejumula’ and established in a three-liter pot: 17 cm diameter and 20 cm height. Media consisted of sterile top forest soil: cow manure: gravel at a ratio of 5:2:1. Plants were grown in the greenhouse at an average temperature of 28º C and watered as needed. After one month when the plants were ~30 cm tall, 20 plants were graft-infected with 5 cm stem scion using side-veneer procedure (Hartmann et al. 1997) on both ‘Ejumula’ and ‘Kakamega’. Table 1 **shows** the different combinations of virus infections with SPLCV, SPFMV, and SPCSV, alone and in possible dual combinations used as treatments in this study. Different treatments were kept in separate insect-proof chambers in the greenhouse to avoid cross-infection.

### Greenhouse multiplication of planting material inoculated with viruses

The different treatments (T1–T16 described in Table 1) were tested at three months after inoculation, by Quantitative Reverse Transcription PCR (qRT-PCR) to confirm the presence/absence of SPFMV, SPCSV and SPLCV. qRT-PCR reactions were carried out as described by Cuellar et al. (2015), for SPFMV the primers 5’-CGC ATA ATC GGT TGT TTG GTT T-3’ and 5’-TTC CTA AGA GGT TAT GTA TAT TTC TAG TAA CAT CAG-3’, and the probe 5’-[6-FAM]-AAC GTC TCC ACG CAA GAA GAG GAT GC-[TAMRA]-3’ were used corresponding to the coat protein region of the genome. For SPLCV the primers 5’-GAG ACA GCT ATC GTG CC-3’ and 5’-GAA ACC GGG ACA TAG CTT CG-3’, and the probe 5’-6FAM-TAC ACT GGG AAT GCT GTC CCA ATT GCT-TAMRA-3’ were used corresponding to ACI fragment of coat protein as described by Ling et al. (2010). Plants that tested positive as expected were rapidly multiplied in seedling trays to generate enough material for field trials. During multiplication, a new sterile scalpel blade was used to cut scions to avoid cross contamination between treatments. To ensure that adequate planting material was available for field experiments, plants with double/multiple viruses were multiplied in extra trays due to slow/stunted growth. The multiplied planting material was further randomly tested by qRT-PCR to confirm their infection status before planting in the field.

### Field experimental design

Field trials were conducted for two seasons at the Kenya Agricultural and Livestock Research Organization, Kiboko Centre, Makueni County in Kenya. The Centre is situated at Latitude S 02 º 12.781’, longitude E 037 º 43.078’ and 931 meters above sea level. The soils were sandy loams for each trial in both seasons. Mean annual rainfall in the region is 50 mm with mean monthly maximum temperature of 33 º C. The two seasons of planting were three months apart. The first field trial was established in September 2017 to February 2018 while season II was set up in December 2017 to May 2018. Both trials were laid using a randomized complete block design (RCBD) with three replicates for the sixteen treatments. The land was ploughed, harrowed and ridges prepared by hand at the two sites before planting. Each replicate (plot) comprised 40 plants at inter and intra-row spacing of 1 m and 0.3 m, respectively. Vine cuttings were four weeks old (~ 30 cm long) at the time of planting. Plants were watered immediately after planting and watered by overhead irrigation for 3 hours at night every four days. Weeding was done manually using hand hoes twice a month in the first two months and once thereafter until the crop was harvested. Two rows of finger millet were planted between each plot to reduce spread of viruses between plots by insect vectors. To monitor whitefly abundance, a yellow sticky card trap (26 cm^2^) was placed horizontally at canopy height at the center of each plot. These were replaced after every two month. To minimize further spread of viruses between plots; plants were sprayed fortnightly by alternating systemic and contact insecticide as described by manufacturer on the container product label.

### Evaluation of SPLCV, SPFMV and SPCSV under field conditions

Disease symptom evaluation was done at 30, 90 and 120 days after planting as described by Hahn et al. (1981). A severity score of 1–5 was used, where 1 = plants showing no symptoms; 2 = virus symptoms just starting to appear and this can be as mild chlorotic spots on the older leaves or mild vein clearing or mild purpling at the leaf margin of mature leaf; 3 = the symptoms in 2 enlarge and become more visible; 4 = infected plants showing severe disease symptoms including leaf purpling, leaf chlorosis and leaf shape starts to get distorted; and 5 = infected plants showing very severe virus disease symptoms including total distortion in leaf shape, stunted growth, mosaic, leaf chlorosis and sometimes complete death of an infected plant. At the end of both growing seasons, cross contamination between plots was evaluated by testing with RT-PCR. Three leaves (third/fifth/seventh) were collected from ten plants in the inner middle rows, placed between filter papers and put in a ziplock bag with silica gel. Silica gel was changed several times when the color changed from blue to pink to ensure that the leaves were well desiccated. Samples were pooled into one per plot and tested for SPCSV, SPFMV (RT-PCR) and SPLCV (PCR) as described above to check if any cross contamination of viruses had occurred between plots.

### Root yield assessment

Plants were harvested at 150 days after planting (DAP); 15 February 2018 and 15 May 2018, respectively. Storage roots were graded as marketable (good-quality roots of 100–1200 g) or unmarketable (<100 g). Sixteen parameters were collected during the experiment. These included: disease severity, main branches length (cm), vine vigor (rate of shoot growth-vine strength, diameter and internode length) - Gruneberg et al. (2010), weight of vines per plot (Kg), number root per plant, number marketable storage roots, number non-marketable storage roots, weight marketable storage roots (Kg), number non-marketable storage roots (Kg), total Root yield (t ha^−1^), marketable root yield (t ha^−1^), foliage yield (t ha^−1^),% of commercial root yield, ratio root length/diameter, root dry matter content (%) and harvest index (HI).

### Statistical analysis

The GLM procedure in SAS (ver. 9.1; SAS Institute Inc., Cary, NC) was used for analysis of variance. The two season data were analyzed and are presented separately and combine means for the two seasons. Separation of means was achieved by Tukey’s Studentized Range Test. In addition, analysis of variance was used to test for differences between treatments and treatment means were separated by Fisher’s protected t-test least significant difference by GenStat (2003). Further, PCA and Pearson correlation coefficients showing pair-wise associations of traits for yield and yield contributing characters was generated by XLSTAT to show the relationship of key parameters measured and treatments.

## Results

### Symptom expression and virus detection in single or mixed infection by SPFMV, SPCSV and SPLCV under field conditions

Analysis of variance for disease severity taken at 90 day after planting showed a significant (*F pr.* <.001) interaction between virus treatments for both ‘Ejumula’ and ‘Kakamega’ for the two seasons as shown in Table 2 and Figure 1. ‘Ejumula’ was more severely affected than ‘Kakamega’ for the different virus treatments. Uninfected control treatments for ‘Ejumula’ and ‘Kakamega’ respectively, did not display symptoms compared to virus-infected treatments. Disease severity scores in both seasons differed among treatments. SPLCV infected plants produced mild symptoms in the two varieties used in this study (Fig 2A & 3A). Plants exhibited slight rugosity and upward curling or rolling of leaves. Disease severity due to SPCSV and SPFMV alone was appreciable in both cultivars and both seasons. Uninfected controls were symptomless (Fig 2B & 3B). Purple rings characterized the symptom expression due to SPFMV (Fig 2C & 3C). SPCSV displayed purpling on older leaves (Fig 2E & 3E). A combination of SPCSV+ SPLCV had a more pronounced severity in both seasons on ‘Ejumula’ - showing chlorotic spots and rugosity; while it was less severe for ‘Kakamega’-showing purpling of older leaves and upward curling (Fig 2D & 3D). SPVD and SPVD+SPLCV were the most severe for both seasons. These included: vein chlorosis, purple spots, mosaic, leaf narrowing, deformation and stunted growth (Fig 2F & 3F). Worth noting was that symptom severity declined at 120 DAP in both seasons. RT-PCR/PCR tests performed from bulk samples at the end of the experiment just before harvest indicated the following plots were contaminated with viruses with which they had not been pre-inoculated: ‘Ejumula’-17 (contaminated by SPLCV), ‘Kakamega’ - 24 (contaminated by SPCSV), Kakamega - 18 (contaminated by SPFMV), in season I and ‘Kakamega’ - 18 (contaminated by SPCSV) in season II (Table 1). Because tests were done on bulks, we were unable to determine the extent of contamination, but considering the overall low level of cross-plot contamination observed in bulk testing we assume it was limited to no more than one or two plants/bulk. We could not ascertain the mechanisms of infection for the specific plots, time of infection and if the infection contributed to yield loss. However, as there was no statistically significant difference with the other replicates for the same treatment and symptoms corresponding to those of the contaminating viruses were also not obvious in these plots, we assume they responded to few and late season infections that had minimal impact on plant performance and this was thus not considered during analysis. As no wild *Ipomoea* or sweetpotato fields were present at or near the field trial plots, the source of virus contamination was most likely from adjacent plots.

**Table 2.**
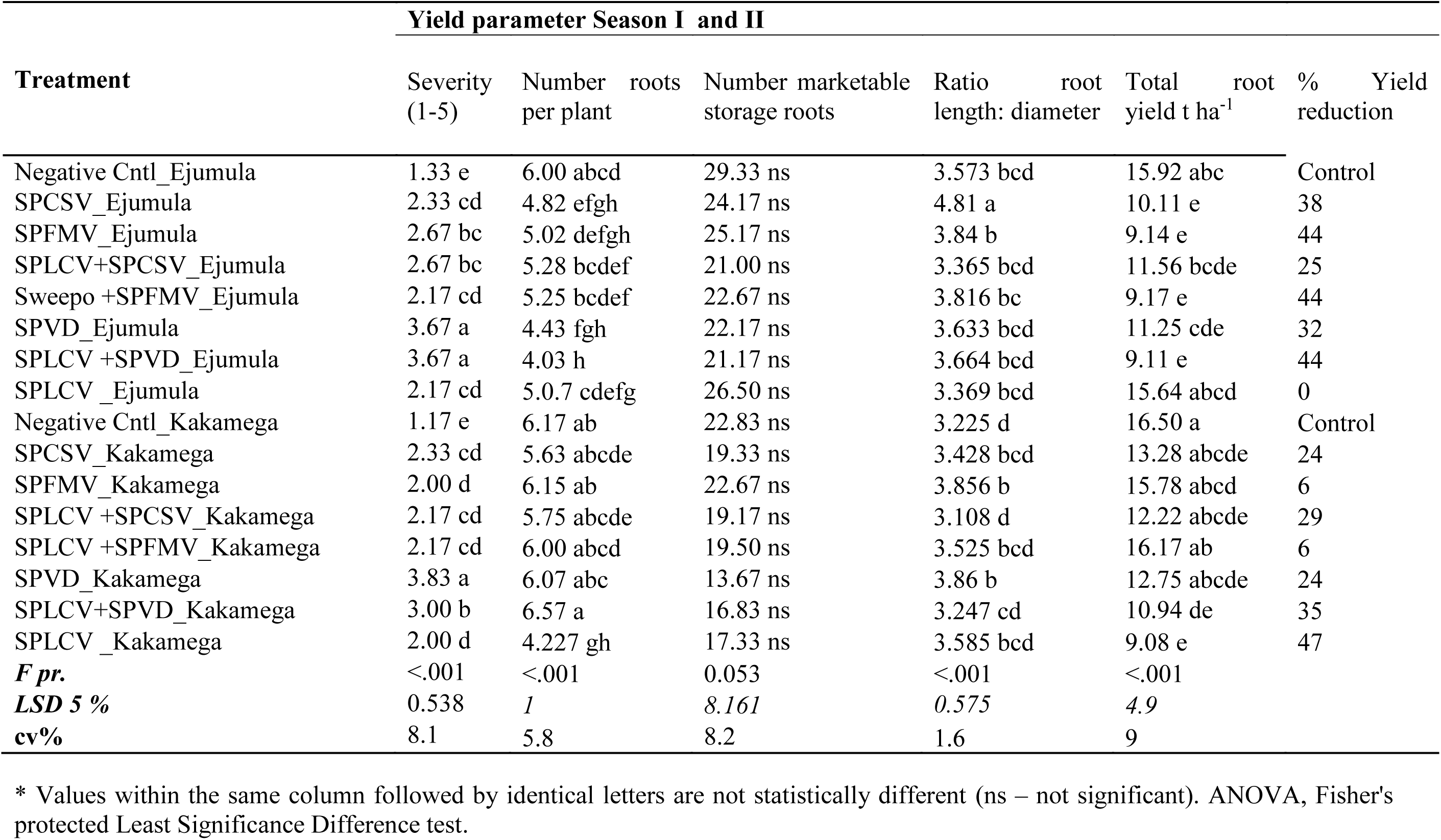
Yield parameters for sweetpotato inoculated with *Sweet potato leaf curl* virus (SPLCV), *Sweet potato feathery mottle virus* (SPFMV), and *Sweet potato chlorotic stunt virus* (SPCSV), alone and in all possible combinations on varieties Ejumula and Kakamega for Season I and Season II combined; Values shown are means for the 16 treatments.

**Figure 1:**
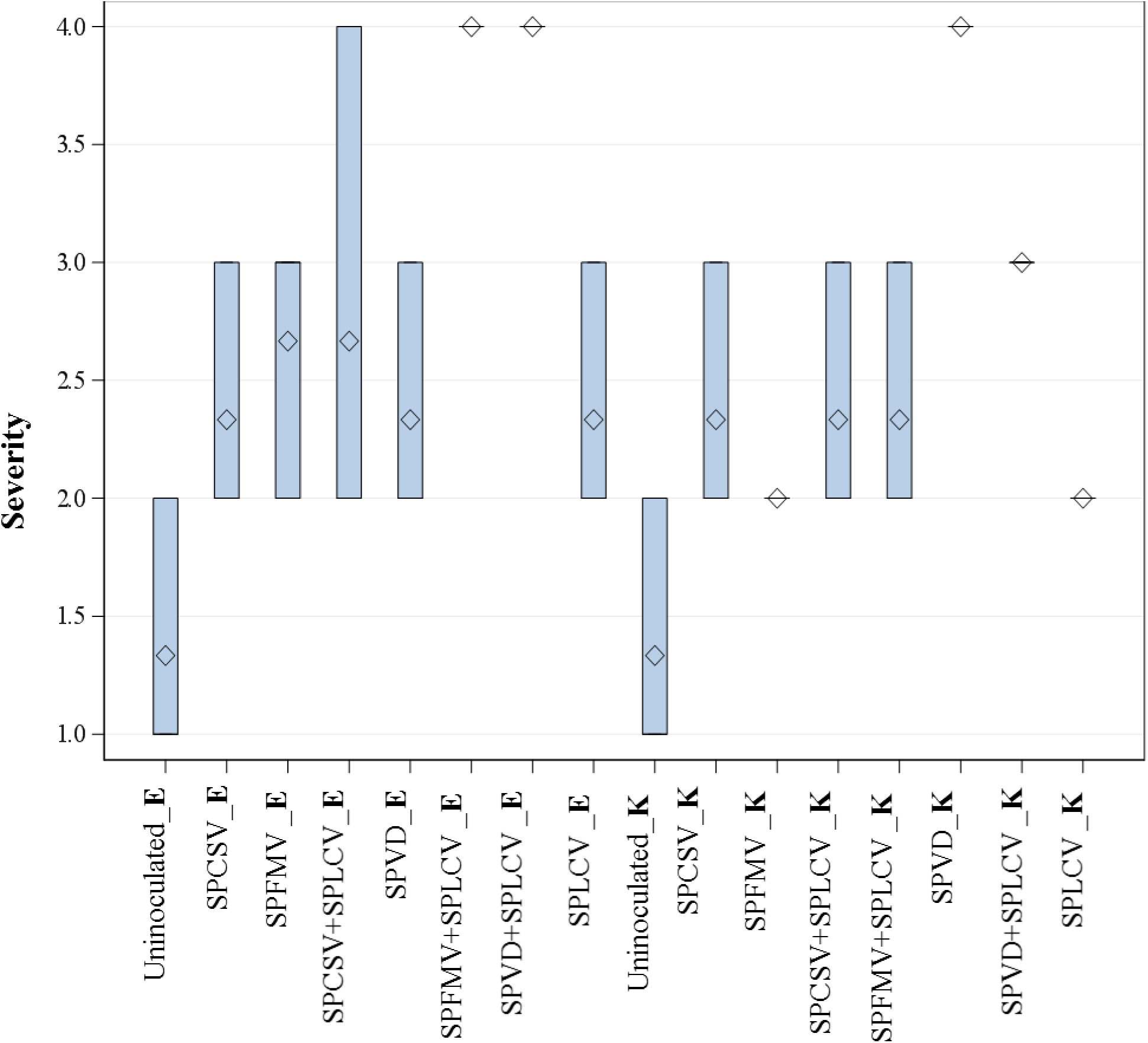
Box-plot for disease severity inoculated with different viruses for combined means for season I_II; expressed varying level of disease symptoms. Severity score of 1 depicts mild symptom expression while 5 is pronounced. All the treatments for Ejumula are abbreviated **E** and **K** for Kakamega.

**Figure 2:**
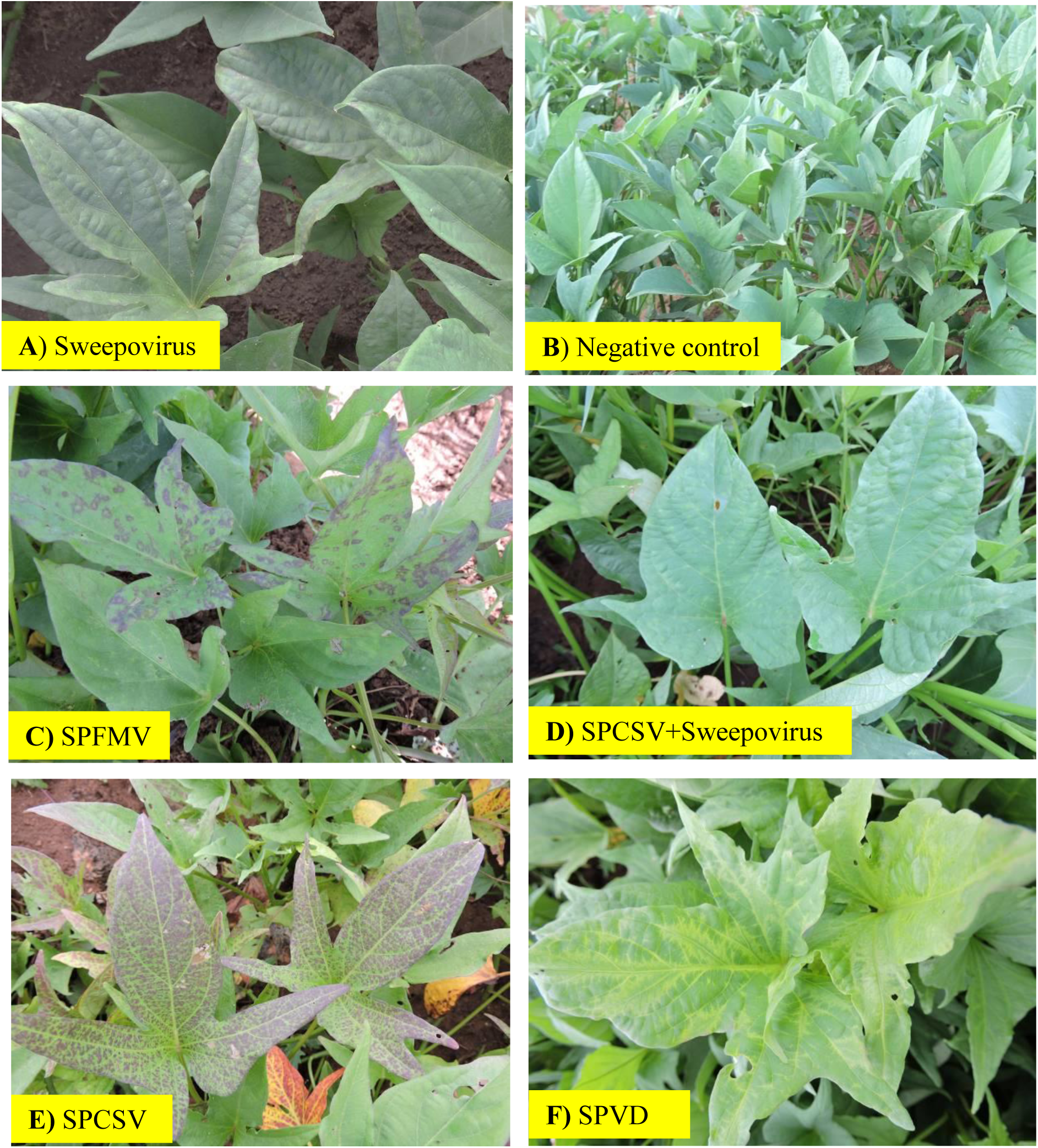
Symptom expression due to single or mixed infection on ‘Ejumula’ by *Sweet potato leaf curl virus* (SPLCV), Sweet *potato feathery mottle virus* (SPFMV), and *Sweet potato chlorotic stunt virus* (SPCSV) under field conditions. A – rugosity due to SPLCV, **B** – uninfected, **C -** purple spot due to SPFMV, **D -** rugosity and chlorotic spots due to SPCSV+Begomo, **E -** purpling of older leaves due to SPCSV and **F -** vein clearing, chlorosis, leaf reduction/deformation–SPVD.

**Figure 3:**
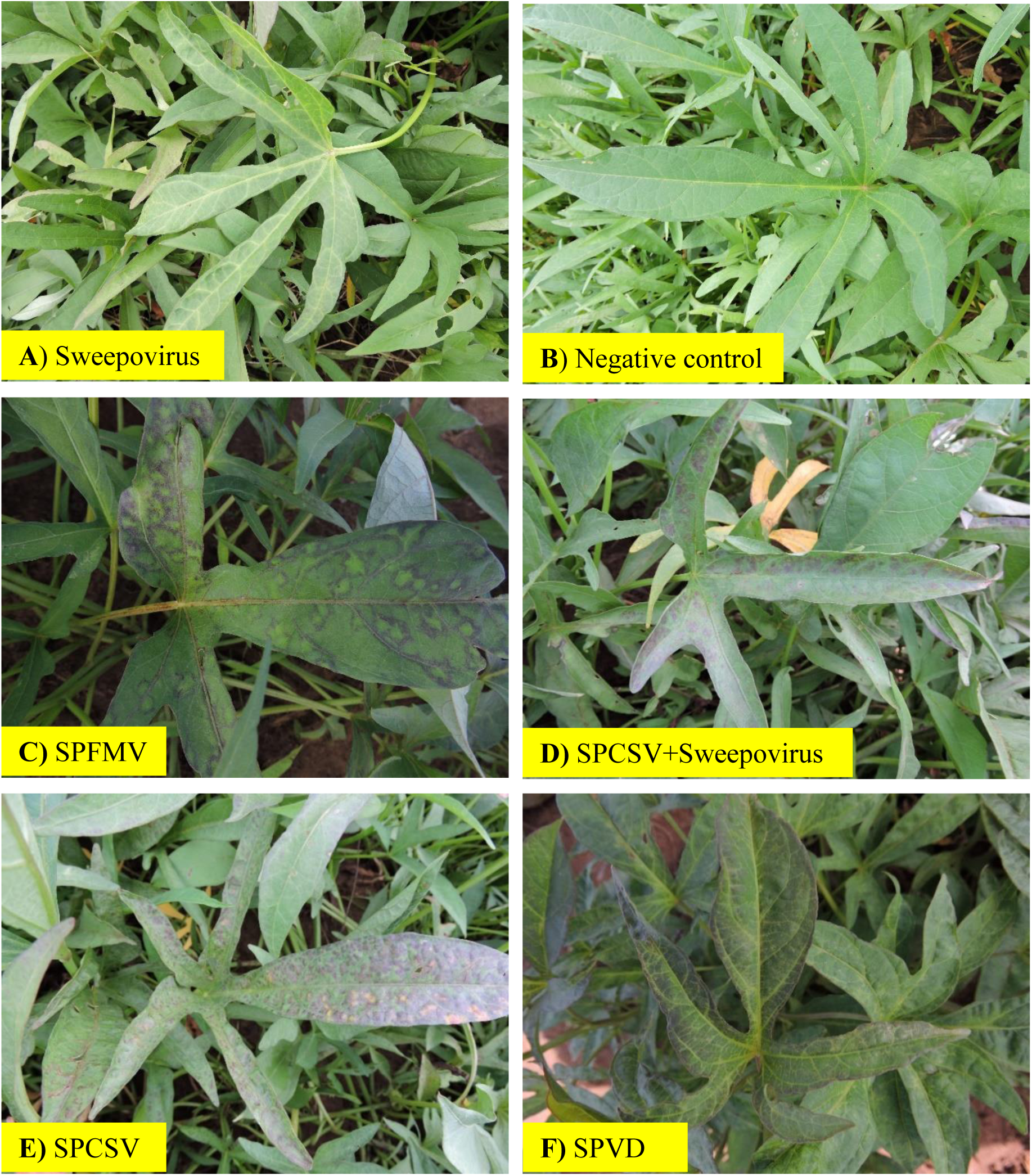
Symptom expression due to single or mixed infection on ‘Kakamega’ by *Sweet potato leaf curl virus* (SPLCV), Sweet *potato feathery mottle virus* (SPFMV), and *Sweet potato chlorotic stunt virus* (SPCSV) under field conditions. A – chlorosis and vein clearing to SPLCV, **B** – uninfected, **C -** purple spot due to SPFMV, **D -** purpling and roll up due to SPCSV+Begomo, **E -** bottom left – purpling of older leaves due to SPCSV and **F -** vein clearing, chlorosis, leaf reduction/deformation–SPVD.

### Effects of virus infection on total root yield

Season I resulted in a greater yield and storage root number (**Supplementary Table 1**) than season II, despite symptoms being generally milder (**Supplementary Table 2**). Significant differences (*F pr.*<.001) were detected, among treatments for root yield related traits (the number of roots per plant, number marketable storage roots, total storage root yield (t ha^−1^) and ratio root length: diameter) Table 2. Analysis demonstrated that total root yield differed significantly for different treatments, variety and season. Uninfected control treatments for ‘Ejumula’ and ‘Kakamega’ respectively gave a higher storage root yield (t ha^−1^) compared to the different virus treatment as shown in Figure 4. ‘Kakamega’ infected with SPLCV or all three viruses had a significant yield reduction of 47% and 35% respectively. ‘Kakamega’ infected with other single or and multiple viruses gave lesser yield reductions ranging from 6% (SPFMV or SPFMV+SPLCV) to 29% (combinations with SPCSV) but were not significant compared to the control. Contrary, there was no yield reduction for ‘Ejumula’ infected with SPLCV alone. However, ‘Ejumula’ infected with all other combinations gave significant yield reduction ranging from 25 – 44 % compared to the uninfected control (Table 2). Significant differences (*F pr.*<.001) in the ratio root length to diameter, the number of non-marketable roots were observed for some of the different virus treatments. A consistent observation was evident from the virus infected treatments with SPLCV (singly or in combination with SPFMV and or with SPCSV) that produced a high number of fibrous roots compared to the uninfected control treatment (Supplementary Figure 1).

**Figure 4:**
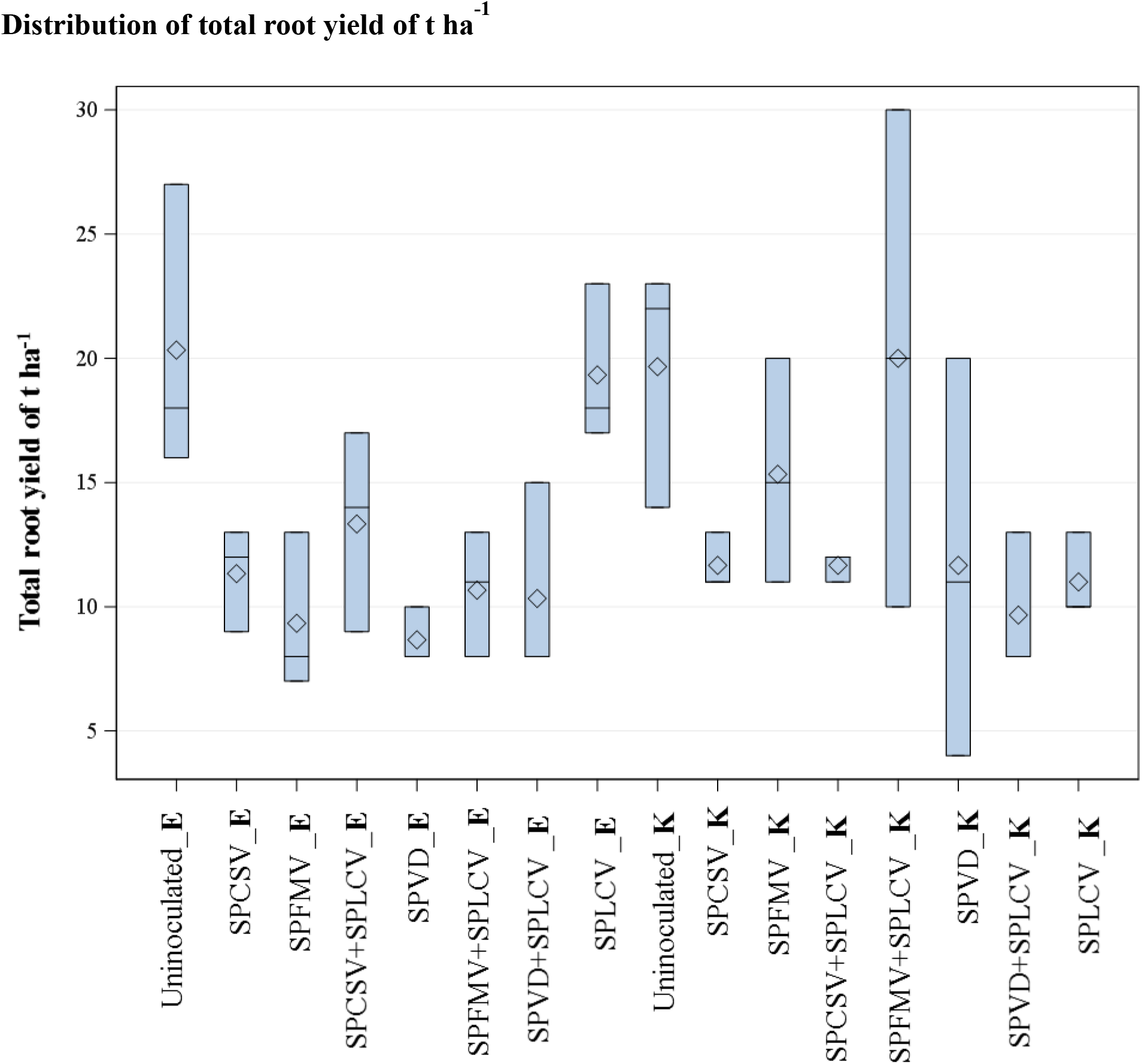
Box-plot root yield in t ha^−1^ for treatments inoculated with different viruses. All the treatments for Ejumula are abbreviated **E** and **K** for Kakamega.

### Yield component traits correlation with the total storage root yield

Yield component traits evaluated for yield and yield contributing characters showed a significant correlation. As observed in Table 3 total storage root yield (t ha^−1^) had strong significant positive association with vine vigor (0.654), marketable storage root yield (t ha^−1^) (0.910), % commercial roots (0.525) and harvest index (0.536). Foliage yield (t ha^−1^) (0.485) and the number of roots per plant (0.338) recorded a relatively strong but non-significant positive association. Contrary, a negative association was observed between total storage root yield and severity at (−0.605). In addition, ratio root length to diameter (−0.387) and the number of non-marketable roots (−0.337) was negatively correlated with severity, though not significant. The results of the correlation biplot (CB) (Figure 5), supports an association of significant correlated traits for yield and yield contributing characters. Sweetpotato yield parameters varied substantially under the virus treatments, and for both varieties. Total storage root yield (t ha^−1^), vine vigor, marketable storage root yield (t ha^−1^), % commercial roots, harvest index, foliage yield (t ha^−1^) and the number of roots per plant displayed furthest away from the center, were most important to distinguish the virus treatments. The PCA, further shows the association of variables (yield parameters) and virus treatments with PCA factor scores in terms of response for the treatments. Uninoculated treatments for ‘Ejumula’ and ‘Kakamega’ had the highest storage root yield (t ha^−1^). Contrary, treatments for ‘Ejumula’ and ‘Kakamega’ respectively infected with SPVD+SPLCV were associated with the highest severity scores.

**Table 3.** Pearson correlation coefficients showing pair-wise associations for yield and yield contributing characters of sweetpotato var. Ejumula and var. Kakamega using virus tested and virus infected (with SPLCV, SPFMV, and SPCSV, alone and in all possible dual combinations) evaluated for two seasons in Kenya.

| Variables | Severity | Vine<br>Vigor | Ratio | Foliage<br>Yield<br>Ton/Ha | No. | Total | Mkt<br>Roots<br>Ton/Ha | % | No. | Harve |
| --- | --- | --- | --- | --- | --- | --- | --- | --- | --- | --- |
|  |  |  | Root<br>Length/<br>Diameter |  | roots<br>per<br>plant | root<br>yield<br>t ha<sup>-1</sup> |  | Comm<br>Roots | Non<br>Mkt<br>Roots | st<br>index<br>(HI) |
| Severity | 1 |  |  |  |  |  |  |  |  |  |
| Vine_Vigour | -0.925 | 1 |  |  |  |  |  |  |  |  |
| Ratio_Root_L_D | 0.204 | -0.189 | 1 |  |  |  |  |  |  |  |
| Foliage_Yield Ton/Ha | -0.485 | 0.364 | 0.180 | 1 |  |  |  |  |  |  |
| No_roots_plant | -0.137 | 0.192 | 0.188 | 0.319 | 1 |  |  |  |  |  |
| Total_Yield Ton/Ha | -0.605 | 0.654 | -0.387 | 0.485 | 0.338 | 1 |  |  |  |  |
| Mkt_Roots Ton/Ha | -0.599 | 0.605 | -0.482 | 0.547 | 0.283 | 0.910 | 1 |  |  |  |
| % Comm_Roots | -0.590 | 0.490 | -0.653 | 0.285 | -0.283 | 0.528 | 0.678 | 1 |  |  |
| No_Non_Mkt_Roots | 0.296 | -0.428 | 0.371 | -0.193 | 0.153 | -0.337 | -0.420 | -0.426 | 1 |  |
| Harvest index (HI) | -0.313 | 0.494 | -0.629 | -0.325 | 0.069 | 0.536 | 0.528 | 0.405 | -0.333 | 1 |

Values in bold are different from 0 with a significance level alpha = 0.05

**Figure 5:**
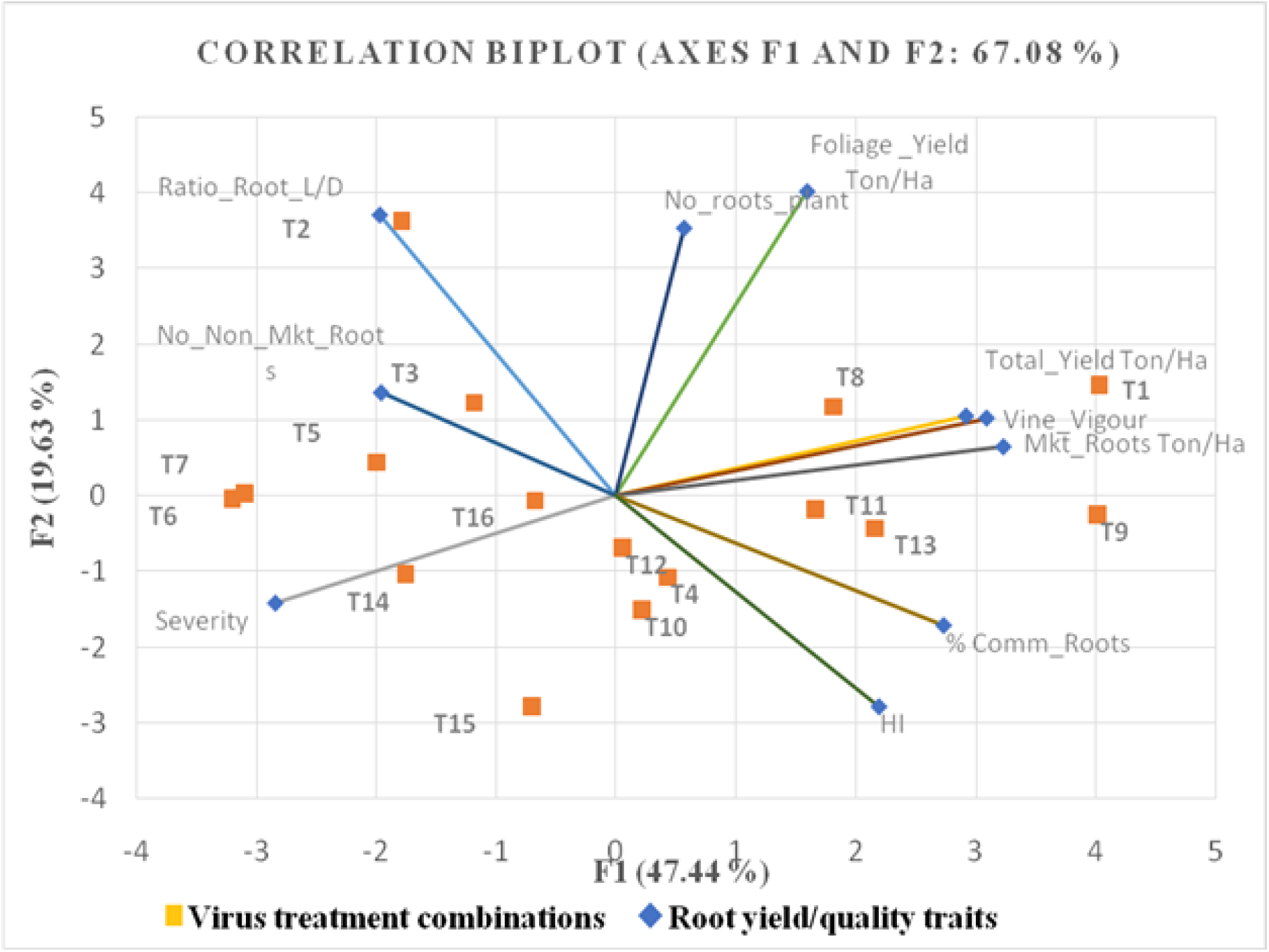
Correlation biplot (CB) representing root yield/quality traits observations and virus treatment variables. Narrow angles depict positively related observations, right angle unrelated and obtuse (wide) angle negatively related. See Table 1 for treatment descriptions.

## Discussion

Sweepoviruses have increasingly been reported from throughout the world. Despite showing few symptoms; increasingly studies are reporting them to have varying but significant impact on root yields. However, until only very recently (Mulabisana et al., 2019), there had been no reports on their impact on African sweetpotato varieties, and this is the first report of their effect on East-African varieties released from local land-races. We show that SPLCV infected plants produced mild symptoms in both varieties used in this study, which however tended to disappear as plants matured. This expands on earlier investigations that reported lack of any symptoms in SPLCV infected plants (Ling et al. 2010; Lotrakul et al. 2003). Similarly, Mulabisana et al. (2019) recently reported mild to no symptoms depending on varieties infected with two different sweepoviruses whereas Cuellar et al. (2015) demonstrated the effect of virus isolates and plant age on symptom expressions and virus titres. Thus, sweepovirus symptomatology can vary depending on cultivar and virus strain and plant age, but invariably is mild and often absent.

Results from our trials show that differences in root yield from SPLCV infected sweetpotato were not significantly different from the uninfected treatments for Ejumula for the two seasons. Contrary, Kakamega had a significant yield reduction following single infection with SPLCV. This illustrates varietal differences in response to sweepovirus infection as found by previous studies, reporting yield reductions between 10-94% between different varieties (Clark and Hoy, 2006, Ling et al. 2010, Mulabisana et al. 2019). Notable findings compared to previous studies were the clear difference in susceptibility to SPLCV between the two cultivars used in our experiments, where ‘Ejumula’ which is relatively susceptible to SPFMV and SPCSV appeared insensitive to SPLCV and ‘Kakamega’ which was more resistant to SPFMV and SPCSV was highly sensitive to SPLCV. This result thus highlights resistance to SPVD and SPLCV (and likely other sweepoviruses) are not necessarily linked. Furthermore, yield losses and symptoms caused by co-infections of SPLCV with SPFMV, SPCSV or both viruses were not significantly different from those caused by the most severe virus in the combination by itself, suggesting a lack of synergistic and limited additive effect of the viruses on yield losses.

SPCSV is considered the most damaging virus of sweetpotato due to its ability to induce synergistic viral diseases with several other viruses (Kim et al. 2017), principal and most severe of which is co-infection with SPFMV, causing SPVD. By itself SPCSV may cause mild to severe symptoms of yellowing or reddening of older leaves, which can often be confused with nutritional deficiencies (Untiveros et al. 2007). Corresponding to the level of resistance of the varieties, single infection by SPCSV induced pronounced symptoms in ‘Ejumula’ (considered susceptible) and produced milder symptoms in ‘Kakamega’ (considered tolerant) and led to yield losses of 38% and 24% on average over both seasons respectively. Co-infection with SPFMV increased symptom severity and yield loss in both cultivars. In ‘Ejumula’ yield losses were significantly different between plants infected by SPFMV, SPCSV or both viruses, whereas in ‘Kakamega’ the yield los between SPCSV, SPFMV, and co-infected plants was identical. This contrasts with most previous reports where more severe yield reductions (from 60-95%) were found when plants co-infected by SPCSV and SPFMV (Milgram et al. 1996; Gibson 1998 and Gutierrez et al. 2003) and may be a result of the specific virus strains and/or varieties used in the current experiment. Further infection of SPLCV in combination with SPFMV and SPCSV, led to slightly higher (non-significant) yield reductions compared to SPFMV and SPCSV alone in both cultivars.

Gutierrez et al. (2003) and Tugume et al. (2013), reported that SPFMV-infected Jonathan and Constanero varieties did not show foliage symptoms under field conditions. Nevertheless, typical symptoms associated with SPFMV were observed in the current study and also by Mulabisana et al. (2019). However, as with SPLCV, a reduction of symptoms was observed in SPFMV infected plants as they matured and most plants became symptomless after 16 weeks. This phenomenon has been reviewed by Gibson and Kreuze (2015) who reported that popular East African cultivars appear to sustain their long-term survival by reverting to symptomless infection and even becoming virus free in some occasions. In our trials, SPFMV by itself had a significant yield impact on ‘Ejumula’, whereas ‘Kakamega’ was not affected, which is in concordance with their level of resistance. Previous investigations have also presented contradictory conclusions regarding yield reductions by single SPFMV infections. Milgram et al. (1996) and, Clark and Hoy (2006) and Gutierrez et al. (2003), noted that single infection with SPFMV did not greatly affect yield. On the other hand, yield reduction of up to 46% were reported in other studies (Gibson et al. 1997; Mukasa 2004; Njeru et al. 2004 and Domola et al. 2008), and recently, Mulabisana et al. (2019) reported reductions of 27-92% by single infection with SPFMV across 12 different cultivars in field trials in South Africa with notable between season effects. Thus, as in the case of sweepoviruses and SPCSV, the impact of SPFMV single infection on yield seem to be highly cultivar specific and across all cultivars globally may be higher than previously assumed. Gibson and Kreuze (2015), have comprehensively documented previous work on yield reductions reported by treatments and cultivars.

Previous investigations have documented that yield and quality of storage roots are sensitive to environmental variations: from year to year, field to field, and even within the same field (Collins et al. 1987; Ngeve and Bouwkamp, 1993 and Bryan et al. 2003). Yield variation among treatments and seasons could be attributed to climatic factors like rainfall and temperature (Roitsch et al. 2003). In season II, we experienced a fourfold increase in rainfall compared to season I (Supplementary Figure 2), but no significant difference in temperatures. Sweetpotato is sensitive to water logging and too much water, specifically early in the growing season, could have led to a lower yield than the first season.

Principal component analysis (PCA) biplot, supports an association of significant correlated traits for yield and help identify yield contributing characters. Furthermore, it shows the relationship of variables (yield and yield contributing traits) and observation (virus treatments) scores, with PCA factor scores in terms of response for the treatments. For instance, uninoculated treatments for var. Ejumula and Kakamega had the highest storage root yield (t ha^−1^). Contrary, treatments inoculated with SPVD+SPLCV for ‘Ejumula’ and ‘Kakamega’ were associated with the highest severity scores and low root yield. Understanding interrelationships among various yield and yield contributing characters is important and can be utilized by breeders when evaluating for virus tolerant varieties during selection.

The highest negative and significant association existed between total storage root yield and disease severity, in both varieties. Gurmu et al. (2015), described a negative correlation between virus symptoms and root yield and is consistent with present results. SPVD is a damaging disease complex of sweetpotato and the negative correlation observed between fresh root yield and disease severity was expected. In addition, ratio root length to diameter, the number of non-marketable roots were negatively corelated to root yield, though not significant. These findings corroborate Bryan et al. (2003) who noted that virus infected planting material produced storage roots with a high length/diameter ratio, culminating in lower total yield and root quality.

## Conclusions

Our study confirmed the relative susceptibility to SPVD of ‘Ejumula’, and revealed it expressed equal sensitivity to both viruses. The relatively SPVD tolerant phenotype of ‘Kakamega’, was expressed as reduced symptoms and absence of yield penalties upon SPFMV infection and reduced symptoms and yield losses upon SPCSV infection as compared to ‘Ejumula’. On the other hand, in contrast to other studies and despite the obvious enhancement of symptoms in SPVD affected plants of both cultivars, we found no evidence of synergistic yield reductions as compared to single infections and suggests that symptoms may not always be an adequate indicator for the effect on yield. This was also clearly the case for SPLCV infection. Considering the widespread presence of begomoviruses globally and also in Africa, this suggests that breeders need to take into account these viruses when selecting for SPVD resistance, as they may inadvertently be selecting for sweepovirus susceptibility by ignoring them. Nevertheless, even in ‘Kakamega’, SPLCV infections induced only mild symptoms that disappeared over time, making such plants difficult to identify to farmers, seed producers and breeders alike to implement any control methods. Thus, adequate diagnostic tests are needed to support these efforts. No effective antisera are available for sweepoviruses and the PCR tests used in this study are too cumbersome for routine implementation in breeding programs or seed certification systems. An effort into developing easier to use molecular diagnostics for sweepoviruses based on isothermal amplifications systems is recommended to support these efforts.

On the other hand, although only one sweepovirus isolate was used in this study, we know from previous studies that this group of viruses is hugely variable and that different isolates differ in their ability to provoke symptoms in sweetpotatoes and indicator plants and accumulate at different titres (Cuellar et al., 2015). Important questions that remain to be answered are if different isolates/species differ in their impact on sweetpotato root yield, if this can be correlated to any particular characteristics other than symptoms (such as virus titres) and if resistance of sweetpotatoes to one of them is correlated with resistance to other isolates. Thus, immediately relevant research topics include evaluating the extent of sweepovirus infections as well as the virus variability in farmers fields in Kenya and Africa in general and the susceptibility to these viruses of current sweetpotato varieties, particularly those selected for resistance to the more visible SPVD.

## Competing interests

The authors declare that they have no competing interests.

## Author Contributions

**Conceptualization:** Jan F. Kreuze, Jan W. Low.

**Data curation:** Bramwel W. Wanjala.

**Formal analysis:** Bramwel W. Wanjala.

**Investigation:** Bramwel W. Wanjala, Elijah M. Ateka, Jan F. Kreuze, Douglas W. Miano.

**Methodology:** Bramwel W. Wanjala, Elijah M. Ateka, Jan F. Kreuze, Douglas W. Miano

**Resources:** Jan W. Low.

**Validation:** Elijah M. Ateka, Jan F. Kreuze, Douglas W. Miano

**Writing - original draft:** Bramwel W. Wanjala.

**Writing - review & editing:** Bramwel W. Wanjala, Elijah M. Ateka, Jan F. Kreuze, Douglas W. Miano and Jan W. Low. All authors read and approved the final manuscript.

## Acknowledgement

The International Potato Center (CIP) funded this study through the Sweet potato Action for Security and Health in Africa (SASHA) project and was undertaken as part of the CGIAR Research Program on Roots, Tubers and Bananas (RTB). The authors specially appreciate the Kenya Plant Health Inspectorate Services–Plant Quarantine and biosecurity Station (KEPHIS-PQBS) Muguga for Laboratory and screenhouse facilities where diagnostic assays and propagation of planting material was conducted. Thanks to the Director General, Kenya Agricultural and Livestock Research Organisation (KALRO) for granting study leave to the first author. We highly appreciate provision of land for undertaking field work. We are grateful to Dr. Daniel Pande for reviewing the draft manuscript. This work is part of a PhD research study by the first author.

---

**Supplementary Figure 1:**
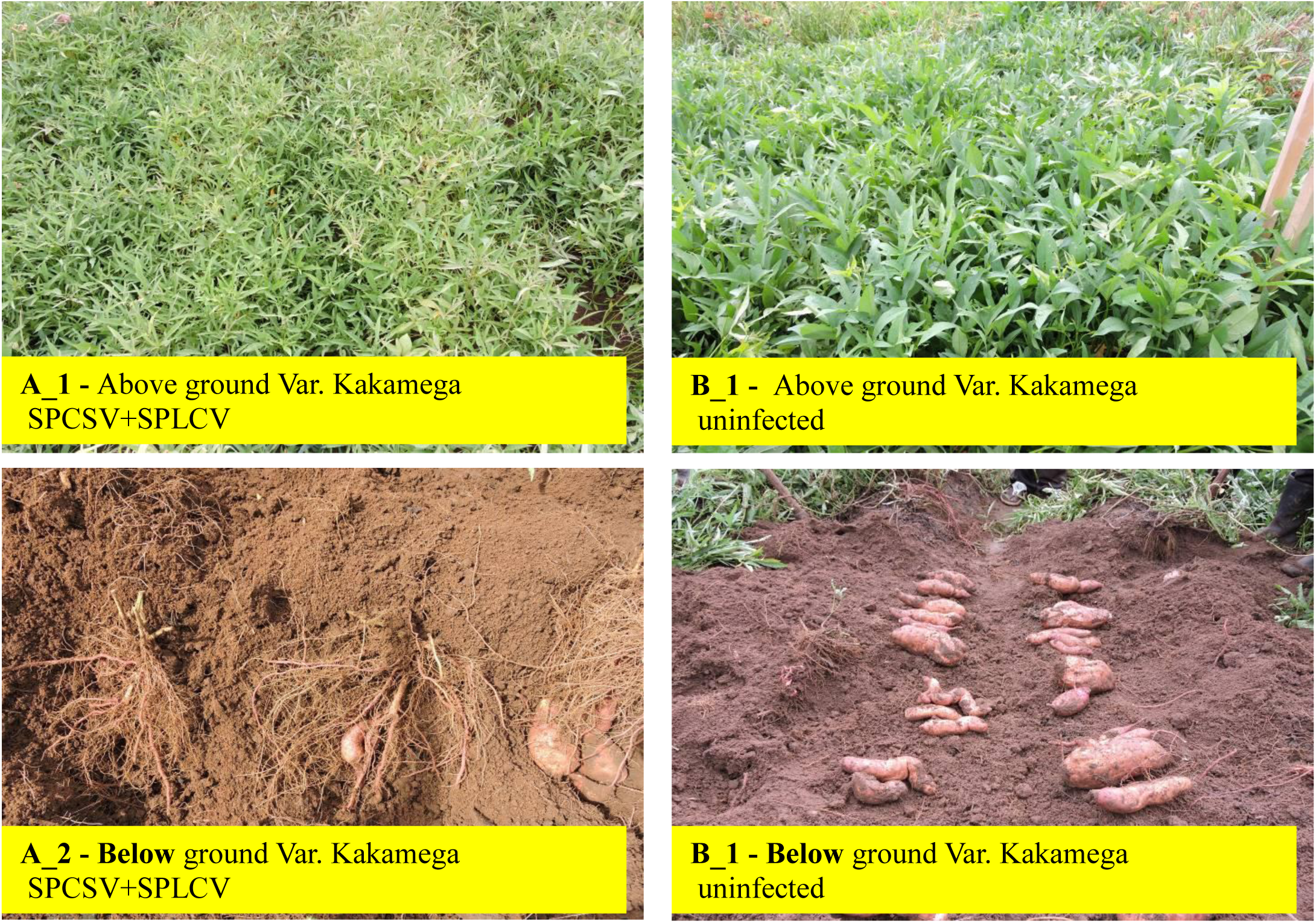
Effect of SPLCV+SPCSV on root formation of Var. Kakamega; A_1 – vigorous above ground cover and A_2 – fibrous root formation and B_1 and B_2 – vigorous ground cover and B_2 – good root formation.

**Supplementary Figure 2.**
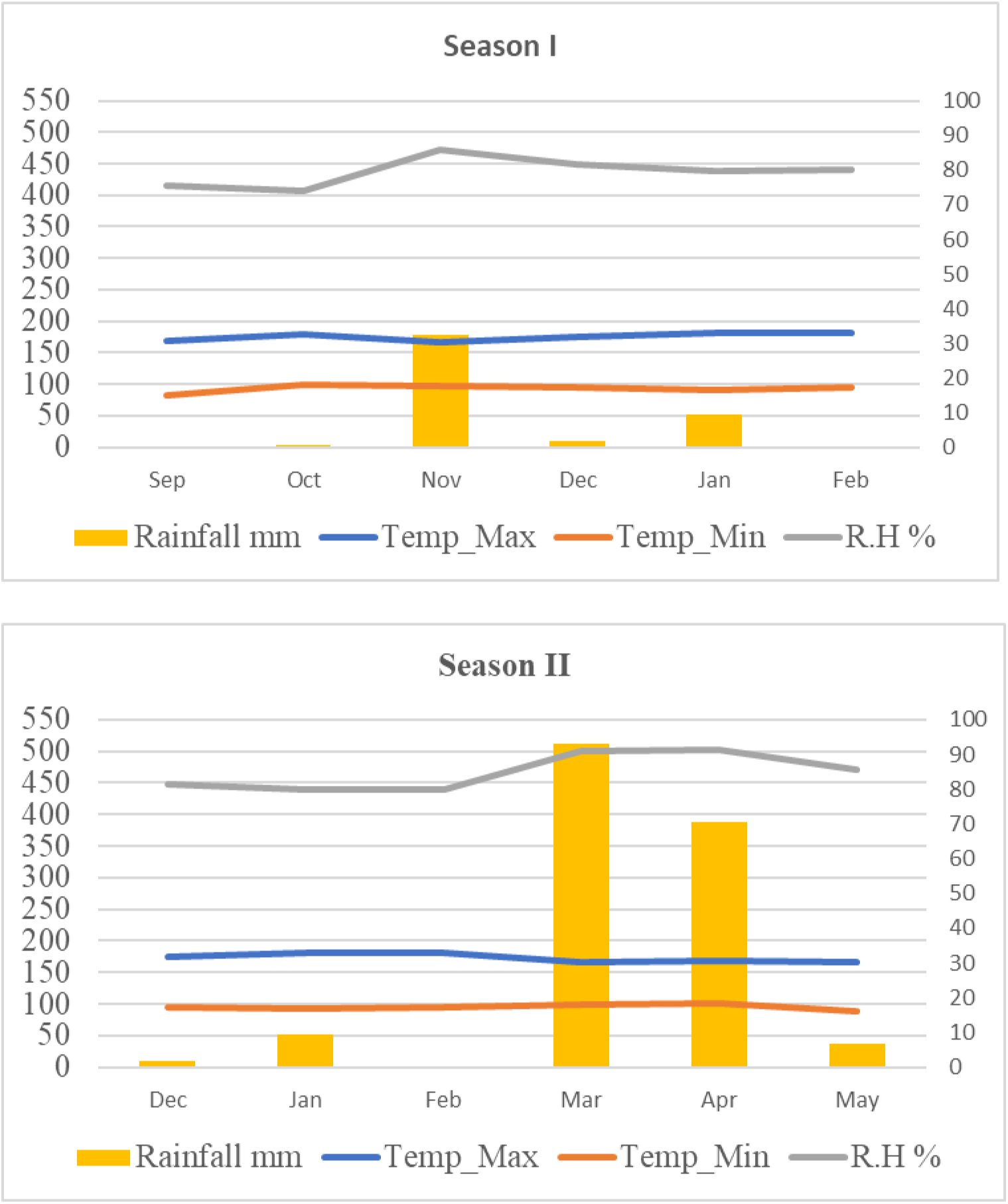
Monthly average climate data (2017 - 2018) rainfall (left axis), temperature, relative humidity (right axis) at KALRO Kiboko, Makueni, Kenya. Data were made available courtesy ICRISAT field station Kiboko.

